# Effect of prescribing and deprescribing oxycodone on pain and function in a mouse model of osteoarthritis: impact of polypharmacy and sex on response

**DOI:** 10.1101/2025.10.22.684045

**Authors:** John Mach, Carina L Blaker, Bryony Winters, Yo Otsu, Elizabeth C Clarke, Heather Allore, Brent Vander Wyk, Kevin Winardi, Roderick Peel, Charlie W Gregson, Neda Assareh, Michelle Kourieh, Sarah N Hilmer

## Abstract

Osteoarthritis and polypharmacy are common in older adults. While not consistent with guidelines, opioid analgesia is commonly prescribed to older adults with osteoarthritis and other causes of chronic non-cancer pain. Long term use of opioids is associated with tolerance, addiction, loss of efficacy and adverse events. Thus, deprescribing (reducing or ceasing) opioids is often required. The effect of polypharmacy on the efficacy and safety of opioid prescribing and deprescribing in this setting is poorly understood. Here we aimed to assess the effects of chronic oxycodone, as monotherapy and in polypharmacy (oxycodone, citalopram, simvastatin, oxybutynin, and metoprolol), and of deprescribing oxycodone, on allodynia, physical and cognitive function, and daily activities in middle-aged, osteoarthritic male and female C57BL/6J mice (n=11-15/ group). Only oxycodone monotherapy reduced mechanical allodynia after 6 weeks of treatment. Chronic polypharmacy with oxycodone transiently increased mechanical allodynia in the injured limb and caused marked reductions in mobility, maximum walking speed, activities of daily living, and increased anxiety. Polypharmacy also altered daily activities detected by an automated animal behaviour recognition cage. Sex differences in treatment response were observed in frailty measurements and activity over 23 hours. Deprescribing oxycodone was well tolerated and did not alter physical or cognitive function but did reverse some of the daily activity changes and mechanical allodynia in polypharmacy treated animals only. This clinically relevant preclinical model provides an opportunity to understand the effects of prescribing and deprescribing opioids in older adults with osteoarthritis, including the impact of comedications and sex.

## Introduction

Opioids are frequently prescribed to older people to treat chronic non-cancer pain, which is commonly due to musculoskeletal conditions such as osteoarthritis (1). Almost half of all opioid prescriptions are for people aged over 65 years (2). However, older people commonly use polypharmacy (5 or more concurrent medications) to manage and treat multimorbidity (3). The effect of polypharmacy on opioid pharmacokinetics, pharmacodynamics, efficacy and safety is poorly understood. Long term use of opioids is associated with tolerance, addiction, loss of efficacy and adverse events (4). In addition, older people are thought to be more vulnerable to adverse opioid effects (e.g., sedation, dizziness, confusion, and falls) because of age-related changes in pharmacokinetics and pharmacodynamics and drug interactions from polypharmacy (5). Thus, opioids are not recommended for long term non-cancer pain management, particularly musculoskeletal pain. However, there is still high chronic opioid utilization globally in older adults, particularly in those with multimorbidity and those with musculoskeletal pain (6, 7).

Deprescribing opioids is a clinical priority which has driven recent development of evidence-based guidelines (8, 9). Deprescribing is the clinical process of ceasing medications where the benefits no longer outweigh the harms, with the goal of improving outcomes (10). Opioid deprescribing can be challenging, even when the potential harms of continuation outweigh the perceived benefits, with an increased risk of overdose and mental health crises observed with cessation of chronic opioids (4, 11). A recent deprescribing study of older people with multimorbidity and polypharmacy reported that of the 286 opioid users at baseline, opioids were discontinued in 58% but 33% of these people restarted opioids (12). While the type of opioid prescribed can predict effectiveness of opioid deprescribing (13), the impact of sociodemographic and clinical factors including polypharmacy on response to opioid deprescribing is unknown.

Preclinical research on polypharmacy can help overcome the limitations of clinical studies in understanding the effects of polypharmacy on opioid efficacy and safety in older age. In healthy animals, chronic polypharmacy that includes an opioid has previously been shown to impair physical function and activities of daily living in both sexes (14) and increase frailty in males (15). Completely deprescribing polypharmacy reversed polypharmacy-induced adverse effects on mobility, activities of daily living and frailty. This indicates removal of all drug burden is beneficial, yet in a clinical setting in patients with multimorbidity, it is unlikely to be feasible to cease all drug treatments. Similarly, opioids are rarely prescribed to patients without pain. It is therefore unclear if deprescription of opioids as a monotherapy or in a polypharmacy regime would show similar benefits in the presence of a painful condition.

Here we use a clinically relevant chronic non-cancer pain model to explore the effects of chronic opioid use as a monotherapy and within a polypharmacy regimen. We investigate the effects of opioid use in monotherapy and polypharmacy, and opioid deprescription, on pain and global functional outcomes (physical function, activities of daily living, frailty, and cognitive function) in male and female mice with osteoarthritis induced in middle-age.

## Methods

### Animals

Male and female C57BL/6J (B6) mice, a standard strain for pharmacologic, toxicologic and ageing research, were investigated in this study. Healthy animals were sourced from Animal Resource Centre, Perth, WA, Australia or the Kearns facility, Kolling Institute, Sydney, Australia, which obtains animals from Animal Resources Centre (ARC). Animals were sourced and aged until the commencement of the study at ∼11 months of age. Mice were housed at the Kearns LAS facility in boxes of no more than two of the same sex with ad libitum access to food and water. A 12-hour light-dark cycle was maintained. This study was approved by the NSLHD (RESP/21/012) and USYD Animal Care Ethics Committee, Sydney, Australia (2022/2045).

### Treatment and disease model details

The study design is shown in figure 1a. From ∼10.5 months, mice were administered standard chow (Rat and mouse premium breeder diet 23% protein, Gordons Specialty Feeds, NSW, Australia) until ∼11 months of age. All mice were then switched to control feed (same chow as medicated feed without medicine). Control feeds contained 20% protein, 4.8% fat, and 59.4% carbohydrate (Standard Meat Free Mouse and Rat Feed, Specialty Feeds, WA, Australia). After 2 weeks of this diet, baseline assessments were performed. For animal well-being, animals were checked and weighed weekly.

**Figure 1.**
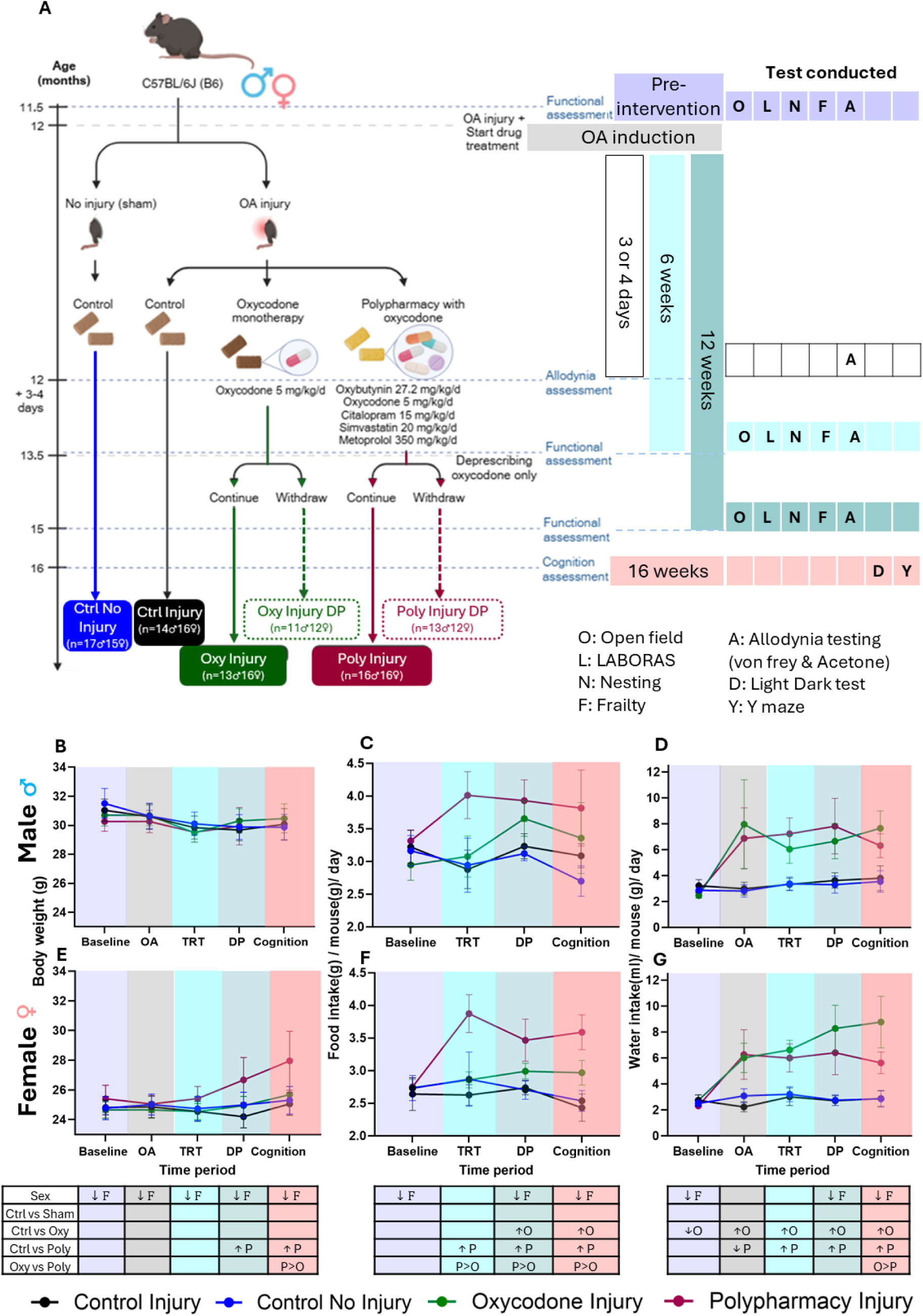
Effect of chronic oxycodone administered as monotherapy and within a polypharmacy regimen on allodynia and functional activity: study design, weight, food intake and water intake. (a) Healthy male and female C57BL/6J (B6) mice aged 12 months were randomized into 6 groups per sex: Control no injury (sham) with no treatment; osteoarthritis (OA) injury with either no treatment (control, Ctrl), oxycodone monotherapy (Oxy), or polypharmacy including oxycodone (Poly). The mice underwent 6 weeks of drug treatment until the age of 13.5 months, when those in the Oxy and Poly groups were re-randomized to either continue treatment or to only deprescribe (DP) oxycodone, over 4 weeks. This study assessed weight and/or function at different time periods; baseline testing (∼11.5 months; purple), OA induction (12 months; light grey), allodynia testing (12 months and 3 or 4 days of age; 3 to 4 days after intervention; white), treatment testing (13.5 months of age; 6 weeks of intervention; cyan), deprescribing testing (15 months of age; 12 weeks of intervention; teal), and the cognition testing point (∼16months of age; 16 weeks of intervention; pink). Test conducted are depicted in table with letters representing experiments and legend below. Sample sizes are displayed. (b, e) line graph displaying body weight, (c, f) food intake (Note that no OA timepoint food intake data is presented, as food intake specific to OA induction could not be accurately extrapolated because data recording occurred at variable days before and after OA induction) and (d, g) water intake over the course of the study for males (top row) and females (bottom row), respectively. Interventions are shown with lines and solid circles; Control no injury (Sham; blue), Control (Ctrl) injury (black), oxycodone (oxy; green) and polypharmacy (Poly; Burgundy). Data displayed as mean ± 95% Confidence intervals. Statistically significant difference (p<0.05) shown in table as indicated by; ↑ higher, ↓lower, O: oxycodone injury, P: polypharmacy injury, M: male, F: female. No sex*treatment interaction was observed. Deprescribed animals were pooled with corresponding treatment groups for the statistical analysis of ‘baseline’, ‘OA’ and ‘TRT’ time points, as deprescribing had not been initiated yet. They were removed from following time points; ‘DP’ and ‘Cognition’. Sample sizes for each test are shown in Supplementary Table 2. Deprescribing body weight, food intake and water intake are shown in Supplementary Figure 2 and Supplementary Table 3.

At 12 months of age, osteoarthritis (OA) was induced unilaterally (under 2% isoflurane 1L/min supplied with oxygen) using a mouse model of anterior cruciate ligament rupture (ACLR)(16, 17). The ACLR involved positioning the knee in a motorised loading apparatus. Increasing the tibial-compressive load caused the femur to slide backwards relative to the tibia, overloading the ACL. A sharp drop in the recorded force confirmed the rupture. The right limb underwent injury, for limb-limb comparison. Naïve control mice underwent anaesthesia without any further intervention.

After injury, sham-injured mice (i.e. those that did not receive the ACLR injury) were maintained on a control (no medications) regimen. In contrast, ACLR-injured mice were randomised into 3 treatment groups: control (no medications), monotherapy opioid (oxycodone), or oxycodone with polypharmacy (with common therapeutic drugs). Oxycodone was administered in the animals’ drinking water with the same therapeutic dose (5 mg/kg/day) in both monotherapy and polypharmacy regimens. In addition to oxycodone administered in drinking water, the polypharmacy regimen consisted of: metoprolol (350 mg/kg/day), simvastatin (20 mg/kg/day), oxybutynin (27.2 mg/kg/day) and citalopram (15 mg/kg/day), which were administered in the chow manufactured by Speciality Feeds (Glen Forrest, WA, Australia). Drug treatments were administered for 6 weeks at which point, mice were approximately 13.5 months of age. After 6 weeks continuous drug treatment, half of the treated animals commenced opioid deprescribing. During opioid deprescribing, the drug was slowly removed by tapering over 4 weeks. Tapering consisted of diluting the oxycodone in the drinking water to administer a half dose of oxycodone for 2 weeks (2.5mg/kg/day), then halving this dose again in the 3rd week (1.25mg/kg/day), before ceasing oxycodone administration completely at week 4 (0mg/kg/day) (Supplementary Table 1). From age 12 months, food intake was measured every second week and water intake was measured weekly to derive drug dose.

### Measured outcomes

The outcomes measured at baseline (12 months old), 6 weeks (13.5 months old) and 12 weeks (15 months old) consisted of allodynia (Von Frey and acetone tests), mobility (open field), activities of daily living (nesting test), frailty (mouse clinical frailty index), anxiety (light dark test) and cognition (Y maze), and daily activities measured with the Laboratory Animal Behavior Observation Registration and Analysis System (LABORAS) automated cage over 23 hours. Outcomes were measured throughout the study, at time points of baseline (2 weeks before OA induction), OA induction (12 months of age), allodynia testing (3-4 days after OA induction for von Frey and acetone test), chronic treatment (after six weeks of treatment) and deprescribing (after six weeks of deprescribing opioids). Cognition was only assessed cross sectionally, four weeks after the completion of the study (16 months old). All animals were acclimatized in the testing room for 30 minutes before all tests except for the LABORAS. The sequence of the battery of tests is shown in figure 1 (supplementary figure 1).

### Allodynia

The von Frey simplified up-down method (SUDO) was used to estimate a paw withdrawal threshold (PWT) an indicator of mechanical allodynia (18, 19). Ten von Frey filaments were used to manually generate reproducible force between 0.02g and 3.92g separated evenly on approximately logarithmic distribution of force. In mesh floor 10×10×10cm cages von Frey filament 6 (0.39g) was used to apply force to the planar surface of the hind paw. The next lower force filament was used upon licking or withdrawal (+), and higher force was used when there was no response (-) until 6 responses (+ or -) were recorded. Two trials were performed on the left hindpaw followed by 2 trials on the right. PWT is calculated from the pattern of 6 responses (19).

The method to measure cold allodynia was adapted from Mitchell, Harley (20). Mice were placed in 10×10×10cm raised mesh floor cages and 20µL of acetone was applied onto the plantar surface of the hind paw using an angled pipette. Paw licking and leg licking (knee region) were counted over a 30 second period, 2 trials on the left hind paw were completed first then 2 trials on the right hind paw.

### Mobility

Mobility was assessed in the open field and LABORAS apparatus. For open field, testing was carried out in red light at approximately 9am-12pm. Animals were placed in the middle of a white plastic box with an open top (50×50×50cm) and activity was recorded over 5 minutes. Distance travelled, immobility time, maximum and mean gait speed were automatically measured using the ANY-maze program (ANY-maze, Stoelting Co.).

### Daily activities

The Laboratory Animal Behaviour Observation, Registration and Analysis System (LABORAS; Release 2.6, Metris, Netherlands) was used to measure daily activities following the conditions specified previously (21). Briefly, animals were placed in the LABORAS apparatus, such that they were caged individually for recording at 10am to 9am the next day (23 hours). The environment was controlled with a regular 12-h light/dark cycle (lights on 7:00 am; off 7:00 pm). Food and drinking water were provided corresponding to their treatment groups. Data was analysed by applying the LABORAS program over hourly segments.

### Activities of daily living

The nesting assessment test was applied to measure activities of daily living as previously described (15). Nesting material (Bed-R’Nest 8G Irradiated (60×25×25mm), Tecniplast Australia Pty Ltd), was introduced to animals from the beginning of the study. To test nesting ability, their houses were removed for 2 days to encourage nest building. After 48 hours, old nesting material was removed, and new nesting material was provided. 24 hours later at 9am-12 noon, nests were scored. Nests were divided into 4 quarters and scored from 0-5 (0: undisturbed, 1: disturbed, 2:flat, 3:cup, 4:incomplete dome, 5:full dome). The score of each quarter was summed to give the total score. JM scored all animals. Animals housed together received the same score.

### Frailty index

The mouse clinical frailty index developed by Whitehead and colleagues was used (22). This test considers 31 different parameters covering clinical deterioration of different systems in ageing mice. Each parameter receives a score 0, 0.5 or 1 and the overall frailty index score is the sum of all parameters divided by the total number of parameters. In this study, frailty testing was performed blinded to treatment groups by JM and two other scientists who were trained by JM (RP and CWG). While frailty index is a continuous measure, animals with a frailty index score above 0.18 are considered frail for analysis (23).

### Anxiety

Anxiety was measured using the light-dark test previously described (24). Briefly the apparatus was an opaque box divided into two chambers (dark and light) with an entry door (7.5×7.5cm). The dark chamber (28 cm wide × 18 cm long × 30 cm high) was a compartment with a lid, and the larger light chamber (28 cm wide × 27 cm long × 30 cm high) was exposed to bright light (65±2 lux). Mice were allowed to explore the light dark apparatus for a total of 5 minutes. They were placed in the light chamber, two-thirds the length away from the entry door to the dark chamber and recorded. Data was analysed with ANYmaze (version 7.09).

The percentage of total session time in the light chamber, compartment transitions and latency for first entrance to dark compartment, were used to assess anxiety levels in mice. Animals that spent less time in the light chamber, had more transitions and shorter latency to first entrance were considered more anxious.

### Cognition

The Y maze was used to measure short-term spatial memory(25). Briefly, in white light, animals were placed in the starting arm and allowed to explore the Y maze with the novel arm closed off for 10 minutes during the training session. After 1 hour, the testing session commenced, where the animal was placed back into the starting arm to explore for 5 minutes with the novel arm open. The exploration latency of the novel arm was measured. Animals with greater memory of the ‘old’ environment are thought to have a natural tendency to preferentially explore new ‘novel’ environments.

## Statistical analysis

For welfare and drug intake, weight, food intake and water intake were accessed at each time point and analysed with general linear models (GLM) adjusted for sex testing treatment effects and generating least squared means (LSM) to comparing treatment groups.

For all outcomes separate sex adjusted GLMs were conducted.

To investigate the impact of injury (control no injury vs control OA injury) the injury main effect was assessed for the change between baseline, and allodynia (3-4 days), treatment (6 weeks) or deprescribing (12 weeks), adjusted for sex using a GLM. To determine whether injury effects differed by sex, a sex*treatment interaction was included and retained if significant and LSM were generated to compare groups.

To assess the treatment effect (control OA injury vs oxycodone injury vs polypharmacy injury) the same GLM model was applied but instead injury main effect was replaced with treatment effect and LSM were generated to compare treatment groups. To determine whether treatment effects differed by sex, a sex*treatment interaction was included and retained if significant.

After the week 6 treatment assessment, the oxycodone and polypharmacy groups were re-randomized to either continue treatment or to only deprescribe (DP) oxycodone over 4 weeks. At week 12 and 16, outcomes were assessed (Figure 1). Among those in active treatment groups, the impact of deprescribing oxycodone compared to original treatment regime was estimated with GLMs adjusted for sex and treatment group. To determine whether deprescribing effects differed by sex, a deprescribing*sex interaction was included and if deprescribing differed by treatment group a deprescribing*treatment interaction was included, these terms were retained if significant, and LSM were generated to compare groups.

Daily activities measured in the LABORAS automated cage included: immobility, locomotion, rearing, climbing, grooming, scratching, eating and drinking, undefined, distance travelled, average locomotion speed, max speed. The data was categorized into periods related to the diurnal cycles (period 1:10-11am, period 2:11am-7pm, period 3: 7pm-12am, period 4:12am-7am, period 5:7am-9am). Separate linear mixed models for each activity used within period hourly change in activity between observations, repeated across the 5 periods and accounting for within animal correlation with a heterogenous first-order autoregressive covariance structure to estimate injury, treatment and deprescribing effects, applying the main effects and interaction effects as defined above in the GLM model.

Statistical analysis was performed using SAS Studio (version 9.4). With each model, to control the false discovery rate (FDR) across all tests, significance levels were adjusted using the Benjamini-Hochberg procedure with a target FDR of 0.05.

## Results

### Osteoarthritis, treatment and deprescribing were well tolerated

To examine the effect of treatment and osteoarthritic injury on wellbeing, we measured body weight and drug intake (Figure 1). Based on the absence of a reduction in body weight, the interventions were well tolerated. Oxycodone administration increased water intake by approximately 3-fold in oxycodone and polypharmacy groups (Figure 1d, g). Only at cognition testing time point, oxycodone intake was higher in oxycodone monotherapy than polypharmacy group. Deprescribing did not alter body weight but a reduction in water intake was observed (Supplementary Figure 2). Similar results were observed with weight prior to the LABORAS testing (Supplementary Figure 3 & 4), however oxycodone monotherapy increased both food and water intake.

### Mechanically induced osteoarthritic injury

We next explored the effects of osteoarthritic injury compared with sham (Supplementary figure 5). After controlling for pre-intervention measurements, injury had no detectable effect on mechanical or cold allodynia, physical function, anxiety or cognition. For the LABORAS automated cage results (Supplementary Table 10), some minor injury effects were observed such as decreased immobility, drinking and increased scratching at one time point. Sexes did not differ in injury response.

### Polypharmacy impairs physical function and causes allodynia, and some effects differed with sex

Chronic medication effects were examined (Figure 2). Oxycodone monotherapy reduced mechanical allodynia (after 6 weeks treatment) and max speed (after 12 weeks of treatment), (Figure 2 a-g). Three days and 12 weeks after OA induction and treatment, polypharmacy increased mechanical pain sensitivity in the right (injured) hind paw. For 6 and 12 weeks after OA induction and treatment, polypharmacy decreased mobility, max speed, nesting ability; and increased immobility time and anxiety (decreased light dark test (LDT) duration in light zone (Figure 2F), with consistent results for compartment transition but no change in latency to dark area (Supplementary figure 6)). Comparing oxycodone monotherapy to polypharmacy, polypharmacy increased right paw mechanical pain sensitivity, decreased mobility and nesting ability, and increased immobility time and anxiety (Figure 2). A sex*treatment interaction was observed for change in frailty score after 6 weeks of treatment, with polypharmacy increasing frailty score only in males (Supplementary figure 7). This interaction was not observed after 12 weeks of treatment. After 16 weeks of treatment, time spent in the novel arm of the Ymaze was not different between the treatment groups, spatial memory was not altered by opioid monotherapy or polypharmacy.

**Figure 2.**
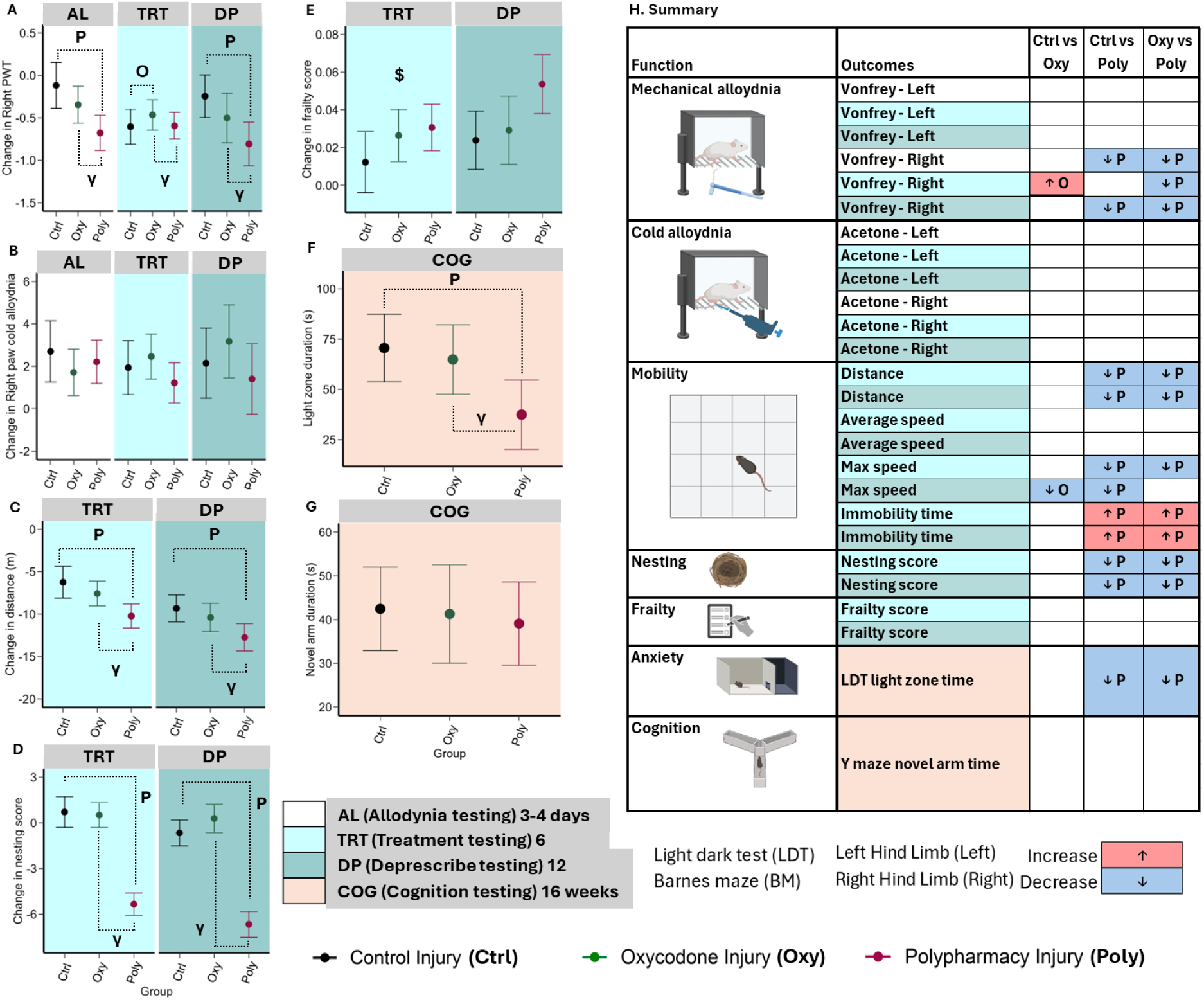
Treatment effect of control (Ctrl; black), oxycodone (oxy; green) and polypharmacy (Poly; Burgundy) on OA injured male and female mice. The following colours represent the time interval at which the change in treatment effect (post – pre) was measured; Allodynia testing (AL; White); 3 or 4 days of treatment and sham/injury, Treatment testing (TRT; Light blue); 6 weeks, Deprescribing testing (DP; turquoise) 12 weeks, Cognition testing (COG; beige): raw data at 16 weeks. Graphs of outcomes, (a) right paw withdrawal threshold (PWT), (b) right paw cold allodynia, (c) distance travelled, (d) nesting score, (e) frailty score, (f) light zone, and (g) Y maze novel arm duration. Data displayed as mean ± 95% Confidence intervals. Significant difference as indicated by p<0.05 via GLM, For all durations; O: Ctrl vs Oxy, P: Ctrl vs Poly, γ: Oxy vs Poly. $: p<0.05 sex*treatment interaction (Supplementary Table 7). Multiple comparisons were adjusted for with Benjamini Hochberg with FDR 0.05. Baseline results shown in Supplementary Table 4. (H) Summary table of outcomes.

As measured using the LABORAS over 23 hours, polypharmacy affected immobility, locomotion, climbing, rearing, grooming, eating, drinking, scratching, distance travelled and max speed (Figure 3-4). Polypharmacy consistently decreased activity (decreased locomotion, climbing, grooming, scratching, distance travelled, max speed and increased immobility) in the second light phase (11-7pm; time periods 2), and decreased climbing, scratching, max speed, and increased immobility in the first dark phase (7pm-12pm; time period 3). However, during the latter dark phase (12-8pm; time period 4), when most groups generally decreased activity, polypharmacy increased activity compared to control injury (locomotion, rearing, distance travelled, max speed) but decreased scratching. These results were quite consistent between 6-and 12-weeks of treatment. Oxycodone monotherapy had limited effects: 6 weeks of treatment decreased locomotion (first light phase, time periods 1-2; 10am-7pm) and scratching (latter dark phase, time period 4). For 12 weeks of treatment, oxycodone monotherapy again decreased scratching (latter dark phase, time period 4; 12-8pm). Treatment*sex interactions were observed such that males and females had different responses to treatment, particularly with scratching, climbing and max speed during at least one time period (Supplementary figure 8).

**Figure 3.**
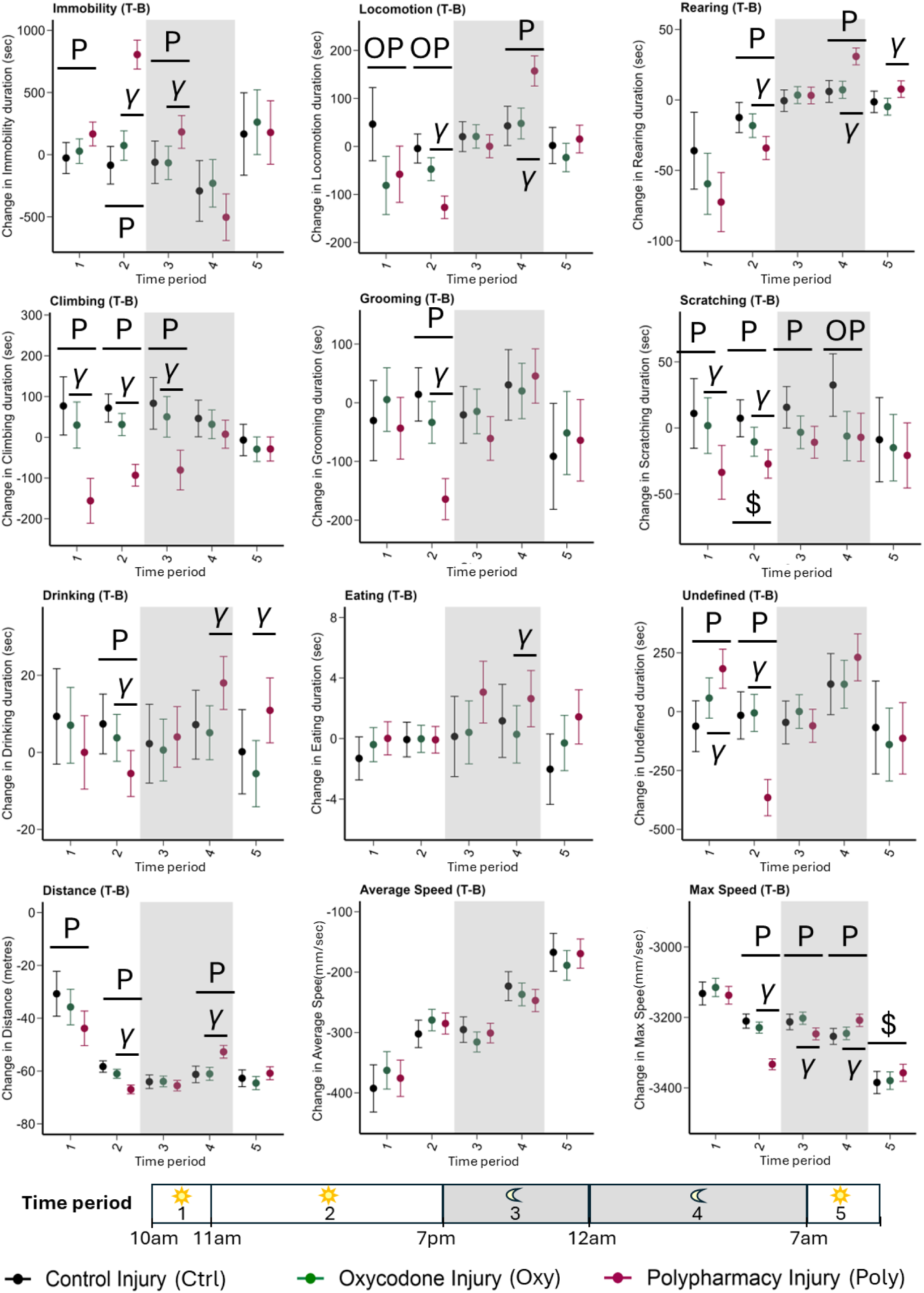
Six-week chronic treatment effect of control (Ctrl; black), oxycodone (oxy; green) and polypharmacy (Poly; Burgundy) on OA injured male and female mice detected using the LABORAS. The graphs represent change in treatment effect over the durations of 6 weeks of treatment (post – pre) for durations of immobility, locomotion, rearing, climbing, grooming, scratching, drinking, eating, undefined and distance travelled, average speed and maximum speed. Different time periods of the 23-hour test; period 1:10-11am, period 2:11am-7pm, period 3: 7pm-12am, period 4:12am-7am, period 5:7am-9am. Shaded areas represent the nighttime (dark). Data displayed as mean ± 95% confidence intervals. Significant difference as indicated by p<0.05 via mixed model with repeated measures applying a first order heterogenous structure, for all durations; **O**: Ctrl vs Oxy, **P**: Ctrl vs Poly, **γ**: Oxy vs Poly, **$**: sex*treatment interaction (shown in Supplementary Figure 8). Multiple comparisons were adjusted with Benjamini Hochberg with FDR 0.05

**Figure 4.**
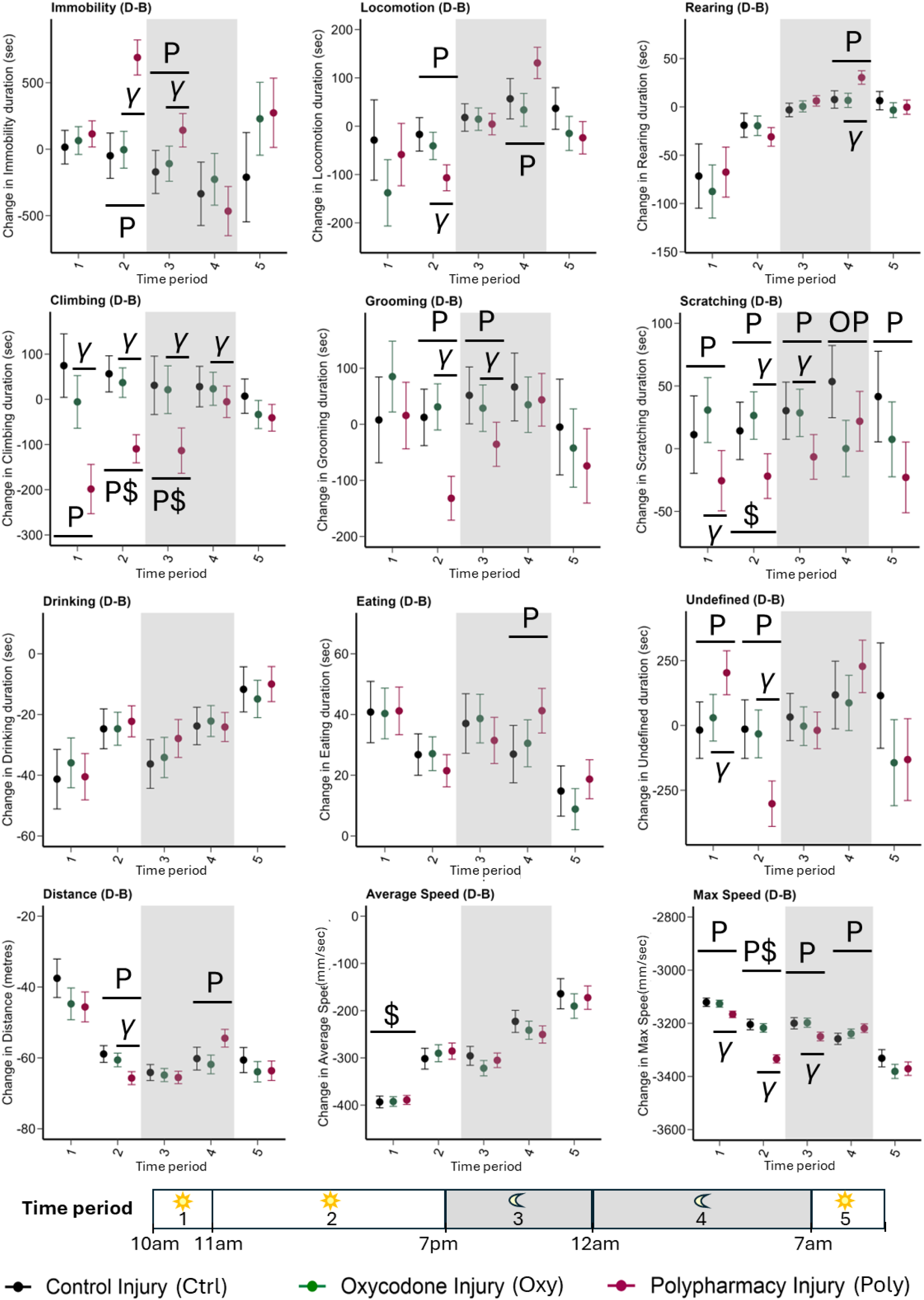
Twelve-week chronic treatment effect of control (Ctrl; black), oxycodone (oxy; green) and polypharmacy (Poly; Burgundy) on OA injured male and female mice detected using the LABORAS. The graphs represent change in treatment effect over the durations of 12 weeks of treatment (post – pre) for durations in immobility, locomotion, rearing, climbing, grooming, scratching, drinking, eating, undefined and distance travelled, average speed and maximum speed. Different time periods of the day; period 1:10-11am , period 2:11am-7pm, period 3: 7pm-12am, period 4:12am-7am, period 5:7am-9am. Shaded areas represent the nighttime (dark). Data displayed as mean ± 95% Confidence intervals. Significant difference as indicated by p<0.05 via mixed model with repeated measures applying a first-order heterogenous structure, for all durations; **O**: Ctrl vs Oxy, **P**: Ctrl vs Poly, γ: Oxy vs Poly, **$**: sex*treatment interaction (shown in Supplementary Figure 8). Multiple comparisons were adjusted with Benjamini Hochberg with FDR 0.05

### Deprescribing oxycodone altered some activities and some effects varied between monotherapy and polypharmacy treatment groups and between sexes

We next assessed deprescription of the oxycodone (Figure 5a-j). Sensitivity analysis comparing no adjustment, adjusting to baseline, and adjusting to change from baseline to treatment commencement time point (treatment - baseline timepoint), demonstrated that ‘no adjustment’ was most sensitive (Supplementary Table 7). For oxycodone monotherapy, deprescribing oxycodone did not change mechanical or cold allodynia. However, for polypharmacy, deprescribing oxycodone reduced mechanical allodynia. Measures of physical and cognitive function were unchanged by oxycodone deprescription. For activities of daily living (nesting experiment), we observed a treatment*sex interaction: deprescribing oxycodone decreased nesting score in females but not males (Supplementary figure 9).

**Figure 5.**
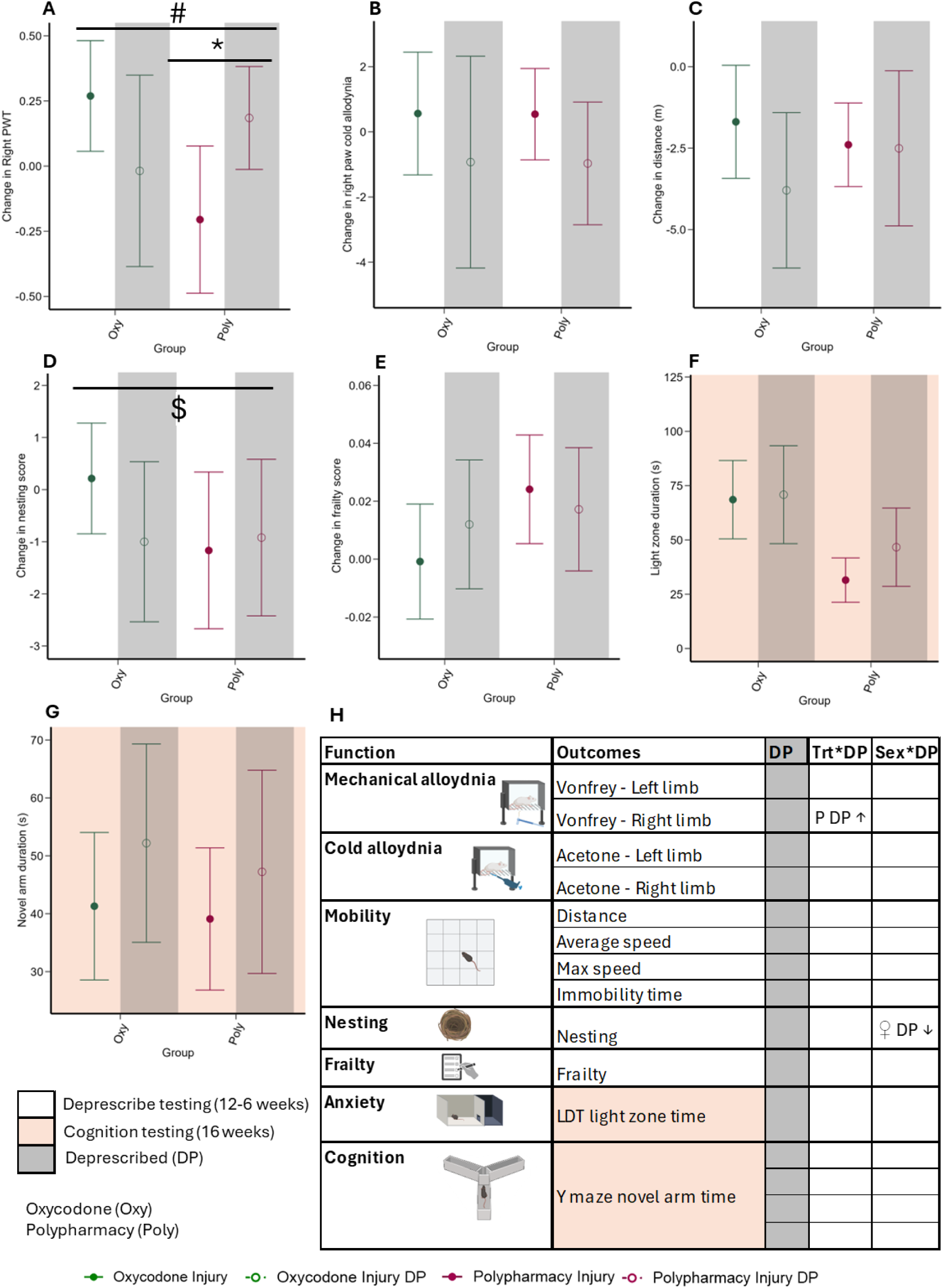
Effects of deprescribing oxycodone on OA injured male and female mice treated with oxycodone (oxy; green) and polypharmacy (Poly; Burgundy). Graphs of outcomes; (A) right paw withdrawal threshold (PWT), (B) right paw cold allodynia, (C) distance travelled, (D) nesting score, (E) frailty score, (F) light dark test light zone duration, and (G) Y maze novel arm duration. (H) Summary table of outcomes. Deprescribing testing (white) compares post (12 weeks) with pre (6 weeks). Anxiety and cognition testing (beige) is cross sectional data obtained after 16 weeks of treatment. DP: Deprescription effect, T: Treatment effect. Deprescribing group is denoted by grey shade and open circle bars. Data displayed as raw mean ± 95% Confidence intervals. Significant difference as indicated by p<0.05, DP: deprescribing main effect, # treatment*deprescribing interaction, and $ sex*deprescribing interaction. Multiple comparisons were adjusted for with Benjamini Hochberg with FDR 0.05 At treatment point before deprescribing, no significant difference was found between continued treatment and deprescribing for both sexes within each treatment group (Supplementary Table 5). Supplementary figure 9 shows sex impact on deprescribing. *p<0.05 as comparing corresponding treatment groups shown by the bar.

Applying the LABORAS (Figure 6), between 11am-7pm (Light, period 2), deprescribing oxycodone increased locomotion, rearing, climbing, scratching; between 12pm-8am (Latter Dark, period 4) deprescribing decreased locomotion, rearing, climbing, max speed and increased immobility; and in the morning 8-10am (Light, period 5), deprescribing decreased locomotion, rearing, distance travelled and max speed, and increased immobility. These effects were observed when deprescribing oxycodone from both oxycodone monotherapy and polypharmacy. Deprescribing effects did differ by treatment group. In polypharmacy treated animals only, deprescribing oxycodone led to a reduction in immobility (Light, period 2), grooming, distance travelled (Latter dark phase, period 4), and grooming (Latter light phase, period 5). In contrast, in the oxycodone monotherapy treatment groups, deprescribing oxycodone resulted in increases in distance travelled (Light, period 2) and drinking (Light and Latter dark phase, period 2 and 4). During Light period 2, deprescribing oxycodone increased average speed for oxycodone monotherapy and decreased average speed for polypharmacy groups. Deprescribing effects on LABORAS activities did not differ with sex (no deprescribing*sex interactions).

**Figure 6.**
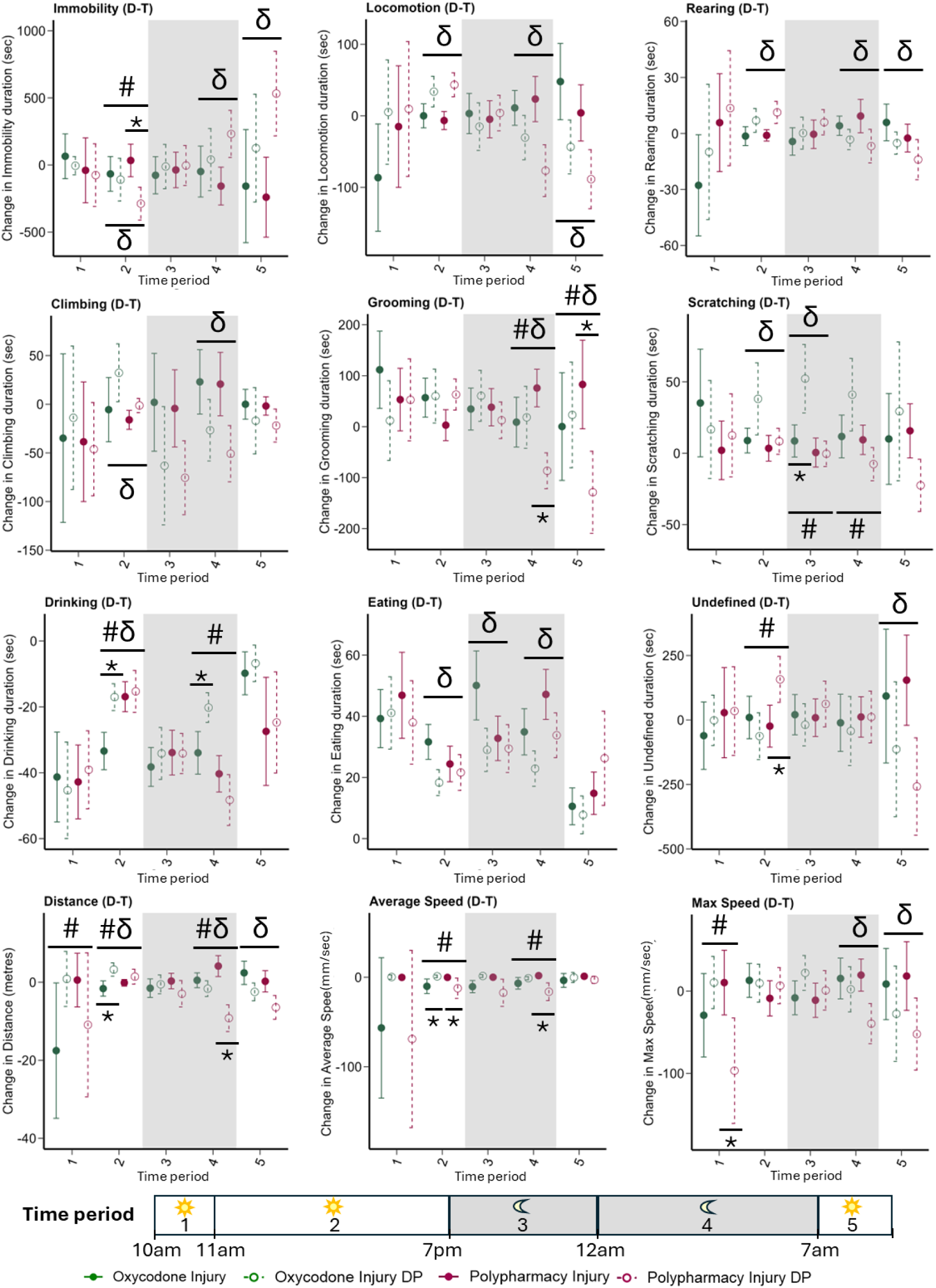
Deprescribing effect (open circle and dotted error bars of corresponding treatment group) on continued treatment oxycodone (oxy; green) and polypharmacy (Poly; Burgundy) in OA injured male and female mice detected using the LABORAS to measure activity over 23 hours. The graphs represent change over the 6 weeks of intervention (post (DP time point) – pre (Treatment time point)) for durations in immobility, locomotion, rearing, climbing, grooming, scratching, drinking, eating, undefined and distance travelled, as well as average speed and maximum speed. Different time periods of the day; period 1:10-11am, period 2:11am-7pm, period 3: 7pm-12am, period 4:12am-7am, period 5:7am-9am. Shaded areas represent the nighttime (dark). Data displayed as mean ± 95% Confidence intervals. Significant difference as indicated by p<0.05 via mixed model with repeated measures applying a first-order heterogenous structure, for corresponding period, δ: deprescribing effect, #: deprescribing*treatment effect. No deprescribing*sex interaction found. *p<0.05 as comparing corresponding treatment groups shown by the bar. Multiple comparisons were adjusted with Benjamini Hochberg with FDR 0.05

## DISCUSSION

In this pre-clinical model of middle-aged onset of osteoarthritis, oxycodone chronic monotherapy had very limited effects, while chronic polypharmacy including oxycodone impaired physical function, activities of daily living, altered daily activities measured using an automatic behavioral recognition cage and increased allodynia, suggesting heightened pain sensitivity. Deprescribing oxycodone reversed some effects of polypharmacy treatment. Some oxycodone treatment and deprescribing effects were impacted by sex and polypharmacy.

The effects of polypharmacy on physical function are consistent with clinical (26, 27) and preclinical studies (28–32). We found that polypharmacy decreased mobility and altered activities of daily living. The underlying mechanisms are emerging, with recent pre-clinical research reporting that polypharmacy causes extensive biologic, molecular changes to a number of organs and biospecimens: brain (28, 29), heart (33), liver (30), drug serum levels (30) and metabolites (31), and microbiome (32).

Some differences between the effects of oxycodone monotherapy and oxycodone in polypharmacy regimens were surprising in the context of the existing literature. There may have been differences in oxycodone exposure between the groups, with monotherapy groups having a higher dose (drinking more water) but drug interactions resulting in higher levels of oxycodone metabolite noroxycodone (30) in the polypharmacy group, or complex pharmacodynamic interactions with the specific polypharmacy regimen used. Clinically long-term use of high doses of opioids is associated with hyperalgesia, which is a paradoxical phenomenon where pain sensitivity increases despite opioid use (34). In the present study, we did not assess for hyperalgesia, however chronic polypharmacy with oxycodone induced mechanical allodynia, after 3 days and 12-weeks of exposure. (30) The additional exposure to citalopram, a Selective Serotonin Reuptake Inhibitor (SSRI), in the polypharmacy group would be expected to reduce pain hypersensitivity(35). Additionally, we find that this regimen of polypharmacy, but not oxycodone monotherapy, caused increased anxiety. Clinically the use of prescription opioids can increase the risk of anxiety and mood outcomes (36), due to neurobiological and chemical changes following prolonged use (37). The decreased activity in the light periods of the LABORAS in the polypharmacy group are consistent with the unadjusted data from a previous study (38) of young and old healthy animals. Further research is needed to determine whether the treatment-specific increased activity in the second dark period (period 4) when animals generally go to sleep, could indicate disturbed sleep, which may heighten pain sensitivity, noting that individuals with disturbed sleep commonly report higher pain sensitivity (39). Interestingly we found that polypharmacy only increased frailty in males following 6 weeks of treatment, demonstrating differences between sexes. Using the LABORAS, we observed many treatment effects, with some responses different between sexes, also highlighting the importance of considering both sexes in pharmacological studies and the need for sensitive systems to detect sex differences in treatment response.

In alignment with clinical studies, this study demonstrated that deprescribing opioids is safe, well-tolerated, did not increase pain sensitivity, effects varied among outcomes, and reversed some of the harms of chronic treatment seen with polypharmacy. In human studies, the effectiveness of interventions aimed at opioid deprescribing and their effects on clinical outcomes vary considerably, with most evidence rated as low certainty (40). Across different interventions, most individuals who reduced or stopped opioid use experienced either improvements or no significant changes in pain and functional outcomes. However, the overall benefits and risks of opioid deprescribing remain unclear due to limited reporting on quality of life and adverse events in the available studies (40). Using a preclinical osteoarthritic model, we observed that opioid deprescription did not reduce body weight, which is a critical measure of animal welfare. While generally, opioid deprescribing did not cause adverse effects, it did decrease nesting ability only in females, which warrants further research. Consistent with clinical observations (41), we found that deprescribing oxycodone had no effects on frailty, mobility (measured using the open field) or cognition. Oxycodone deprescribing did reverse mechanical allodynia and some treatment effects measured by the LABORAS. A previous study exploring this polypharmacy regimen found that complete deprescribing over 6 weeks reversed chronic treatment effects on frailty, mobility, activities of daily living (15) and it reversed some hepatic proteome changes (30). However in the present study, where only oxycodone was deprescribed with the other four medications continued, the medication-related harm observed in the polypharmacy group is likely not limited to the use of oxycodone (42). The finding that deprescribing can reverse treatment effects measured by the LABORAS, suggests sensitive tests to precisely detect subtle changes are necessary to detect benefits from deprescribing, and this is currently being applied to clinical studies (43). Interestingly, following deprescribing of oxycodone from the polypharmacy group, we observed a reversal of the increased locomotion, grooming and distance travelled during the second dark period (period 4), which we hypothesized were due to sundowning syndrome or disturbed sleep. Importantly, we observed that deprescribing oxycodone can have different effects with oxycodone monotherapy and polypharmacy. Future studies are required to validate these findings in humans and explore whether tailored oxycodone deprescribing protocols are necessary for people with differing overall medication use.

Some limitations could be addressed by future research. First, this study did not detect any major injury effect. Osteoarthritis was induced via a tibial compression injury to ensure that all targeted animals developed progressive end-stage structural osteoarthritis (i.e. severe disease). This was done to minimise the variability and uncertainty of spontaneous age-associated osteoarthritis development which has a previously reported incidence of 19% at 15.5 months old and 39-61% from 17 months old (44), with relatively mild degenerative joint disease up to 20 months of age (45). In the present study, we did not observe any effect on mechanical pain sensitisation. This contrasts with changes previously reported in this non-invasive osteoarthritis-inducing injury model in younger 12-week-old animals (46). In two studies of a different surgically induced osteoarthritis model (the destabilizing meniscal injury model) in young adult mice (10-12 weeks of age), one did not observe changes in activity on the LABORAS at 2 or 4 weeks post-surgery, but did see reduced mobility 8 weeks after surgery (47) ,while another reported no change in LABORAS at 12 weeks after surgery (48). One explanation for the absence of change in pain or function at 6 or 12 weeks with induced osteoarthritis in our study is that the middle-aged animals may have already developed spontaneous, osteoarthritis-associated joint and behavioural changes prior to the start of the study. An ageing study of spontaneous osteoarthritic change (45) observed increased mechanical allodynia in both female and male C57Bl/6J mice by 6 months of age although this was only significant in females. By 20 months of age there was evidence of mild structural osteoarthritis in the knee, but notable symptomatic progression occurred in both female and male mice with significantly increased mechanical allodynia, knee hyperalgesia and reduced grip strength compared with young 10-week-old animals. Based on these findings, the similarities between the sham and OA control mice in the present study may be expected and suggest that the structural joint disease severity does not correspond to the severity of behavioural changes, reflecting the disconnect between pain sensitisation and radiographic disease severity observed in osteoarthritis patients (49). Future studies could use microCT and histology or gait analysis to further investigate osteoarthritis severity and disease phenotypes in these animals. Secondly, this study did not investigate the use of paracetamol or non-steroidal anti-inflammatory drugs which are commonly co-administered with oxycodone in clinical practice. Future studies are required to explore varying monotherapy, and combination therapy regimens with different classes and doses of medicines to adequately represent the medication use in older people with osteoarthritis.

Thirdly, the deprescribing effect was investigated only post deprescribing and in the future more frequent measurements should be conducted to better understand the short-term changes (i.e. withdrawal symptoms) following deprescribing to inform and optimise clinical practice. Fourth, drug levels were not measured to confirm whether polypharmacy induced allodynia was due to increased serum oxycodone levels as a result of pharmacokinetic changes. Future studies are needed to investigate biological changes that may provide a mechanistic understanding and consequently inform optimal medication use and deprescribing practice.

The strength of this study is that it provides a potentially translatable model of geriatric pharmacotherapy (50): it investigates chronic exposure to medications that are used by older people in a clinically relevant model of osteoarthritis, a condition commonly causing chronic non-cancer pain in middle and old age. This is the first preclinical study to investigate the effects of deprescribing opioids in the context of polypharmacy. Male and female mice were studied. A comprehensive battery of tests commonly used in pharmacology and aging studies were applied. These methods are comparable to measurements used in humans, such as measuring mechanical allodynia, frailty, mobility, activities of daily living and spatial memory. In addition, automated mouse cage recognition devices, with limited human interference, captured circadian changes. Adequate follow-up was included in this study to assess the effectiveness and outcomes of opioid deprescribing, which is generally a limitation in clinical deprescribing studies that experience challenges with recruitment and retention of participants, as well as limited power to study clinical outcomes (40). The ages at which osteoarthritis-inducing injury, treatment (12 months, middle age) and deprescribing began (13.5 months, still middle age), as well as treatment duration, were selected because they are estimates of clinical observations in humans.

## CONCLUSION

In conclusion, in this study we demonstrate that a polypharmacy regimen that includes oxycodone induces allodynia, increases frailty, impairs nesting ability, mobility and alters daily activities in middle-aged osteoarthritic C57BL/6 mice of both sexes. Sex differences in treatment response were observed in frailty measurements and activity over 23 hours. Deprescribing of opioids was well tolerated, could not reverse treatment effect on frailty, nesting or mobility but could reverse some activities during the day and mechanical allodynia in polypharmacy treated animals only. Importantly, opioid deprescription can have varying effects with monotherapy and polypharmacy treatment. Opioid monotherapy exposure and deprescribing had very little effect on pain sensitisation or function. This model provides an opportunity to understand the effect of polypharmacy and deprescribing of opioids on individuals with osteoarthritis. Future studies can explore mechanisms by which polypharmacy regimens that include opioid induce functional impairment, how deprescribing opioids can reverse some of these outcomes and how sex and treatment regimen (monotherapy or polypharmacy) can impact on the response to opioid treatment and deprescribing.

## AUTHOR CONTRIBUTIONS

S.N.H and J.M conceptualized and conceived the project. S.N.H., J.M., B.W., C.B., Y.O., E.C. designed the study. J.M, K.W., C.G., R.P., conducted majority of the behavioural data and animal welfare check with contributions from B.W., C.B. and Y.O. C.B. conducted all the osteoarthritis induction. NA designed, conducted and analysed majority of the light dark (LD) experiments and guided the interpretation the LD data. H.A. and B.V.W. provided the statistical support. J.M. conducted the statistical analysis and wrote the original draft of the paper with oversight and direction from S.N.H. All authors provided the critical feedback and/or edited the manuscript and approved the final version for submission.

## Acknowledgments

The authors acknowledge the support of the Kearns LAS facility staff, Kolling Institute. The authors acknowledge the technical assistance of Jim Matthews and Dr Alex Shaw of the Sydney Informatics Hub, a Core Research Facility of the University of Sydney. The authors acknowledge expert advice on the allodynia testing and osteoarthritis model from Professor Chris Little and Professor Chris Vaughan. The graphical abstract and experimental design image (Figure 1) were created with https://www.BioRender.com/. Open access publishing facilitated by The University of Sydney, as part of the Wiley - The University of Sydney agreement via the Council of Australian University Librarians.

## List of Abbreviations

ACLR: Anterior cruciate ligament rupture
AL: Alloydnia
ARC: Australian Resource Centre
COG: Cognition
CTRL: Control
DP: Deprescribed
FDR: False Discovery Rate
LABORAS: Laboratory Animal Behavior Observation Registration and Analysis System
OA: Osteoarthritis
OXY: Oxycodone
POLY: Polypharmacy
PWT: Paw Withdrawal Threshold
SEM: Standard Error of Mean
SSRI: Selective Serotonin Reuptake Inhibitor
SUDO: Simplified Up-Down method
TRT: Treatment

---

**Supplementary Figure 1.**
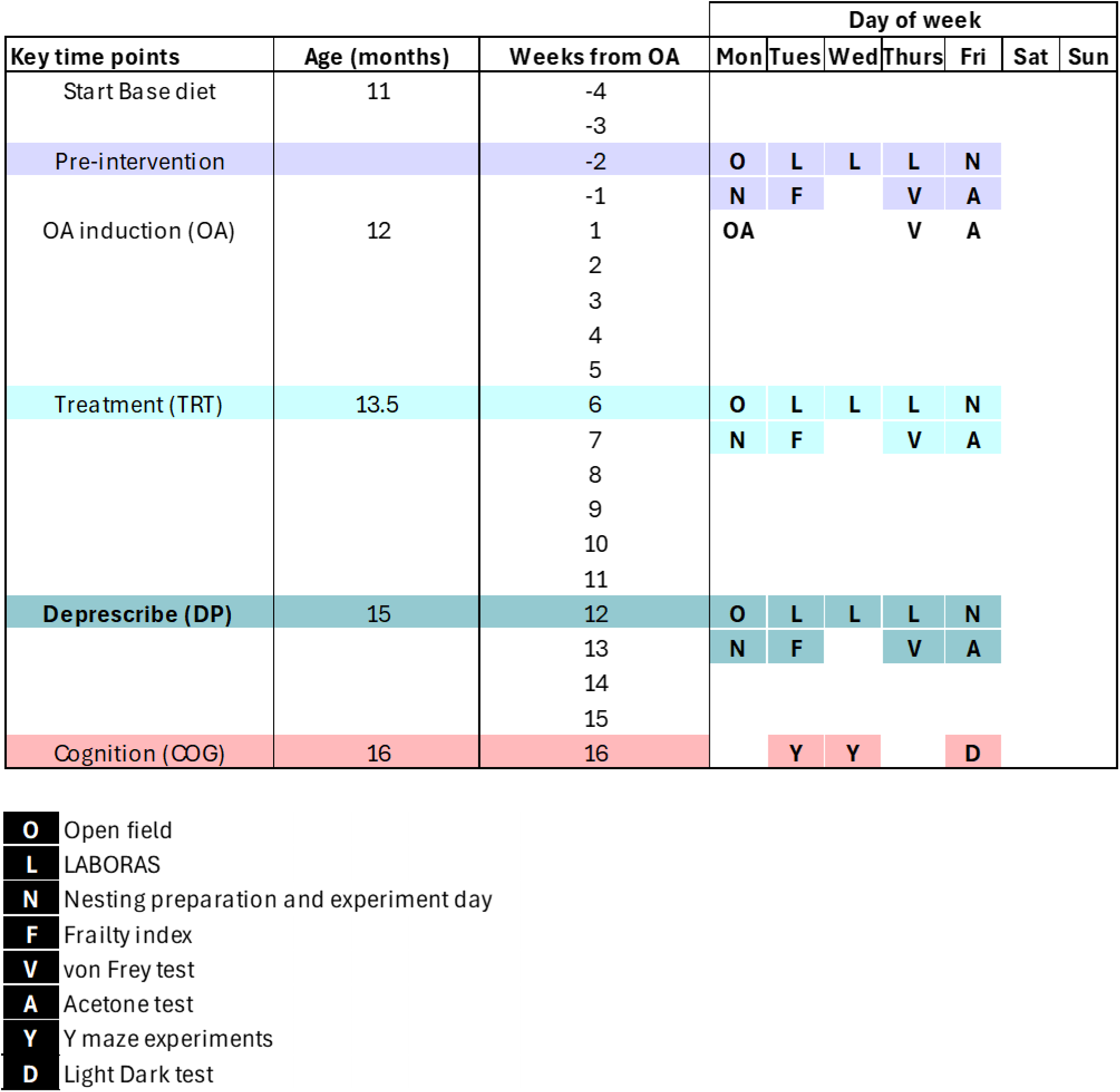
Shows approximate testing of battery of behavior tests.

**Supplementary figure 2.**
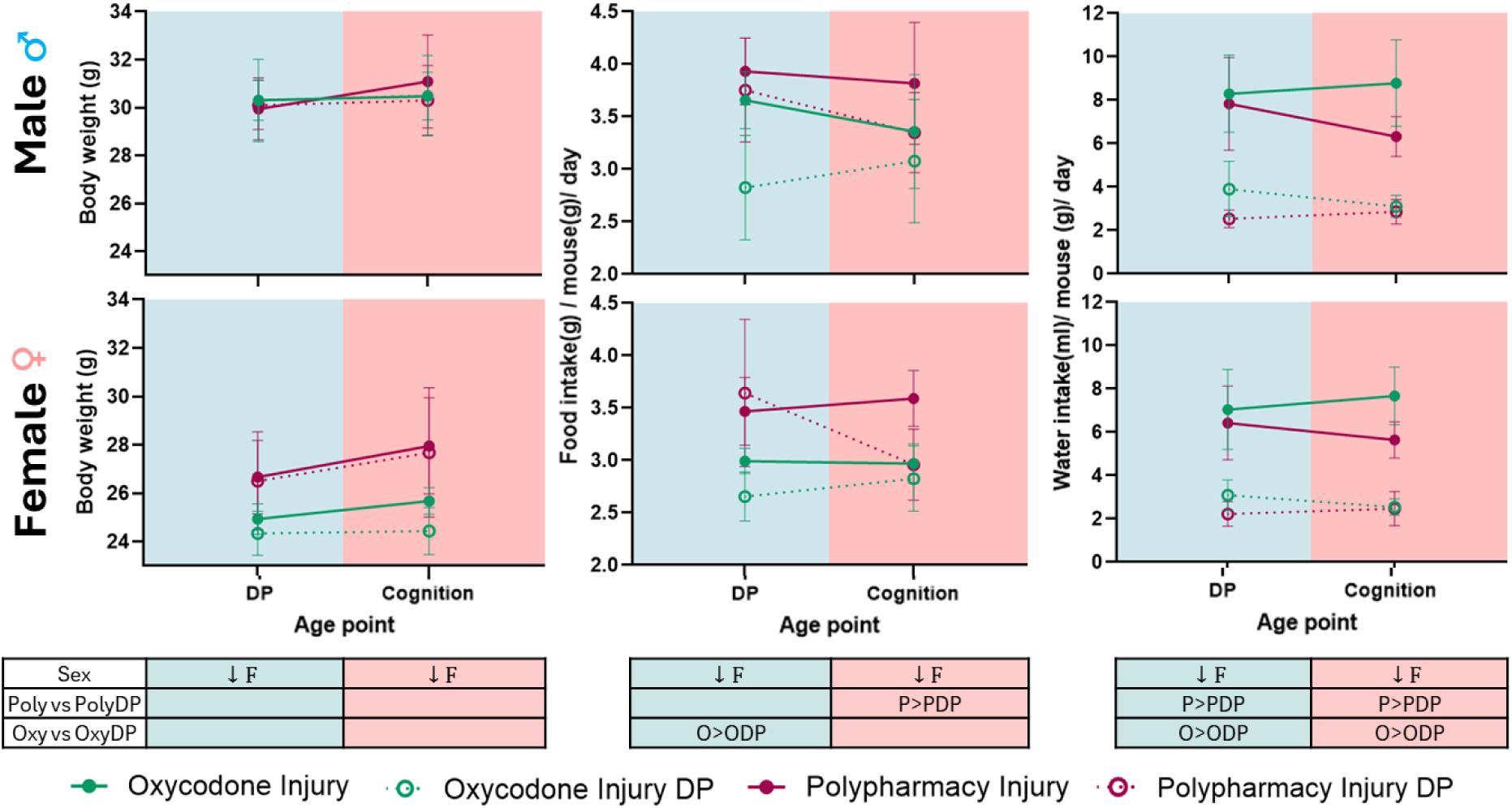
Deprescribing effect on body weight, food intake and water intake. Continued treatment is displayed by solid lines with solid circles; Oxycodone (oxy; green), polypharmacy (Poly; Burgundy), and corresponding deprescribed (DP) groups are shown with the same colour but dotted lines and open circle. Data displayed as mean ± 95% confidence intervals. Statistically significant difference (p<0.05) shown in table as indicated by; ↑ higher, ↓lower, O: oxycodone injury, P: polypharmacy injury, DP: deprescribed, M: male, F: female. No sex*treatment interaction was observed. Only treated and deprescribed animals were included in analysis.

**Supplementary Figure 3.**
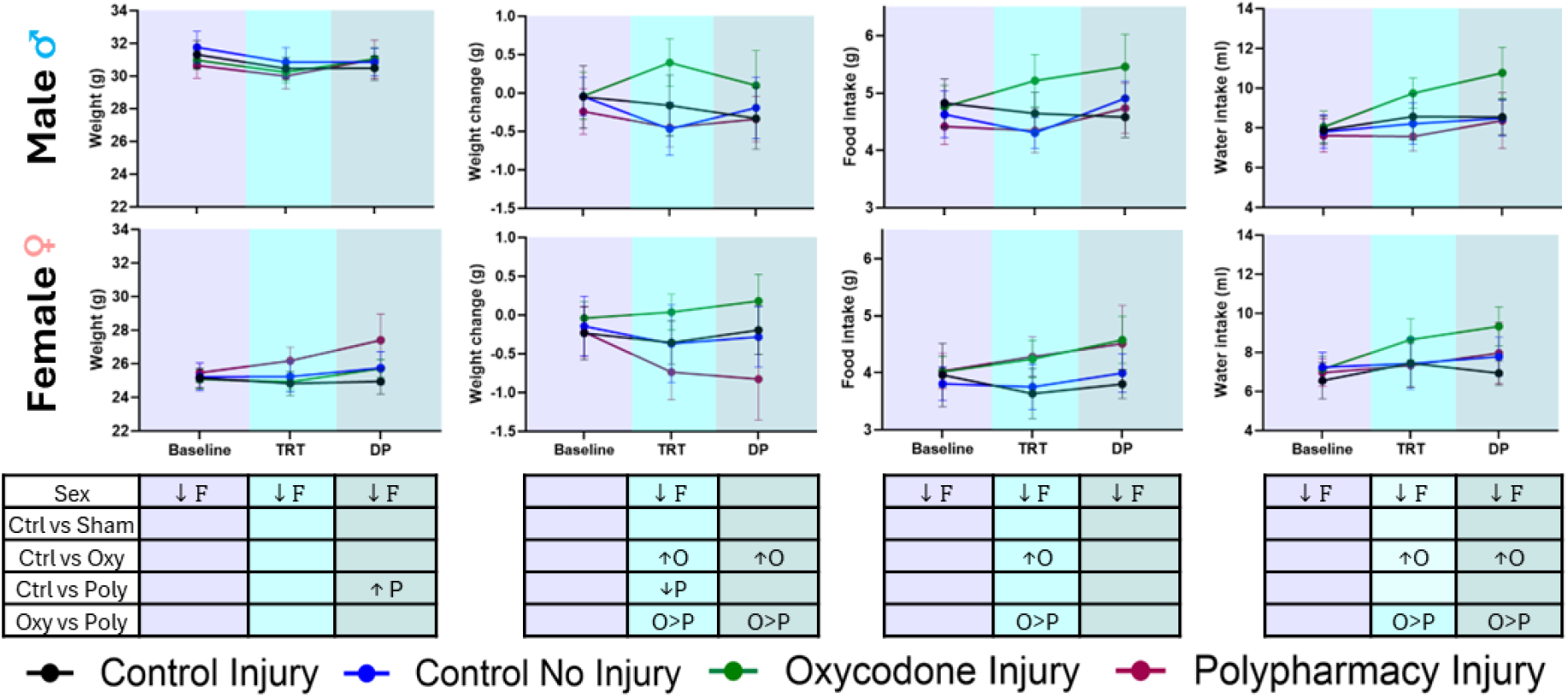
LABORAS experiment, body weight, weight change, food intake and water intake at baseline, treatment (TRT) and deprescribing (DP) time point for males (top row of graphs) and females (bottom row of graphs). Interventions are shown with lines and solid circles; Control no injury (Sham; blue), Control (Ctrl) injury (black), oxycodone (oxy; green) and polypharmacy (Poly; Burgundy). Data displayed as mean ± 95% Confidence intervals. Statistically significant difference (p<0.05) shown in table as indicated by; ↑ higher, ↓lower, O: oxycodone injury, P: polypharmacy injury, M: male, F: female. No sex*treatment interaction was observed. Deprescribed animals were pooled with corresponding treatment groups for ‘baseline’ and ‘TRT’ statistical analysis, as deprescribing had not been initiated yet. They were removed from the DP time point. For LABORAS deprescribing body weight, food intake and water intake are shown in supplementary figure 4.

**Supplementary Figure 4.**
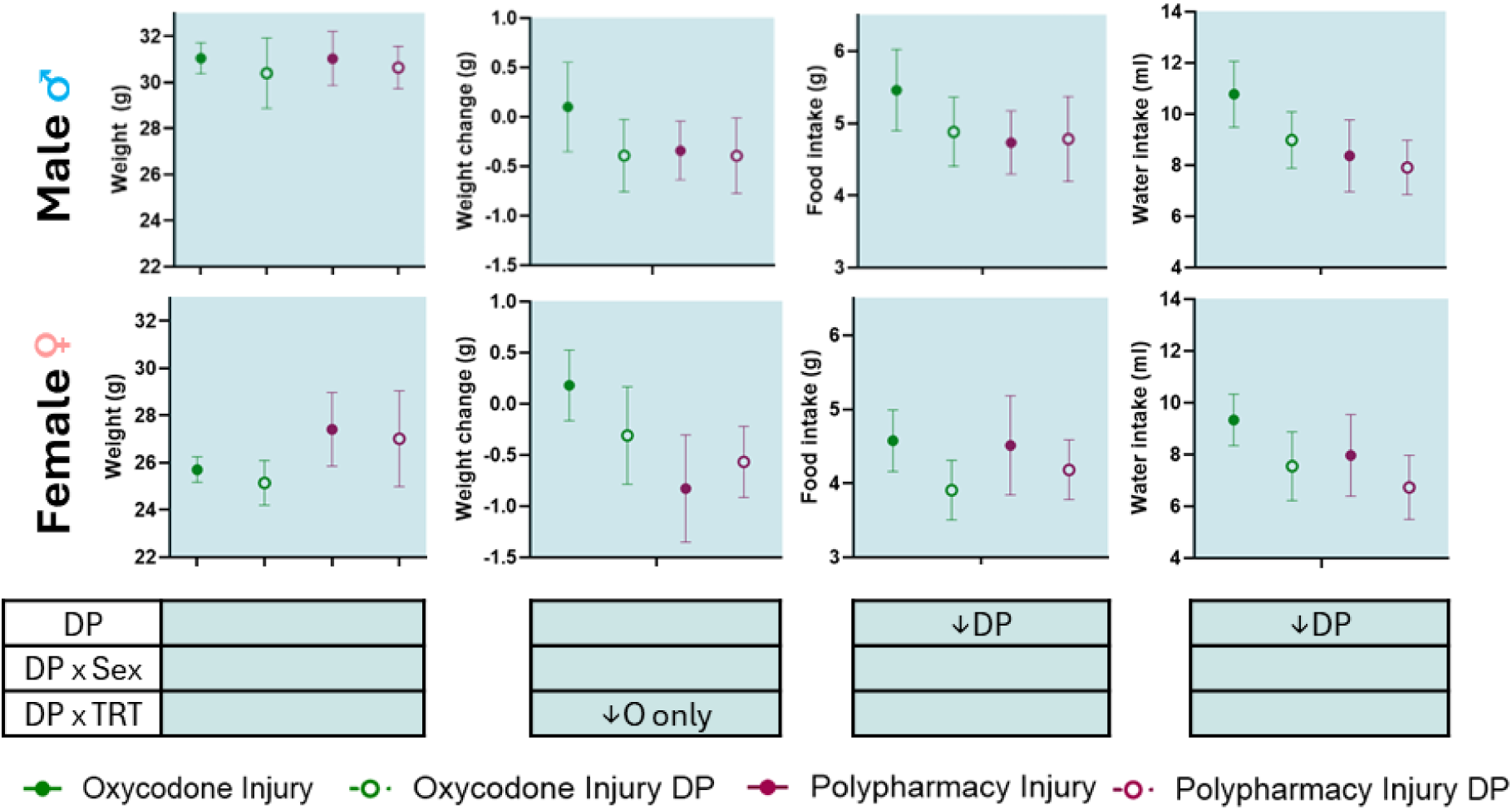
LABORAS experiment, deprescribing effect on body weight, weight change, food intake and water intake at deprescribing time point. Continued treatment is displayed by solid circle; Oxycodone (O; green), polypharmacy (P; Burgundy), and corresponding deprescribed (DP) groups are shown with the same colour but open circle. Data displayed as mean ± 95% Confidence intervals. Statistically significant difference (p<0.05) shown in table as indicated by; ↑ higher, ↓lower, DP: deprescribed. Only treated and deprescribed animals were included in analysis.

**Supplementary Figure 5.**
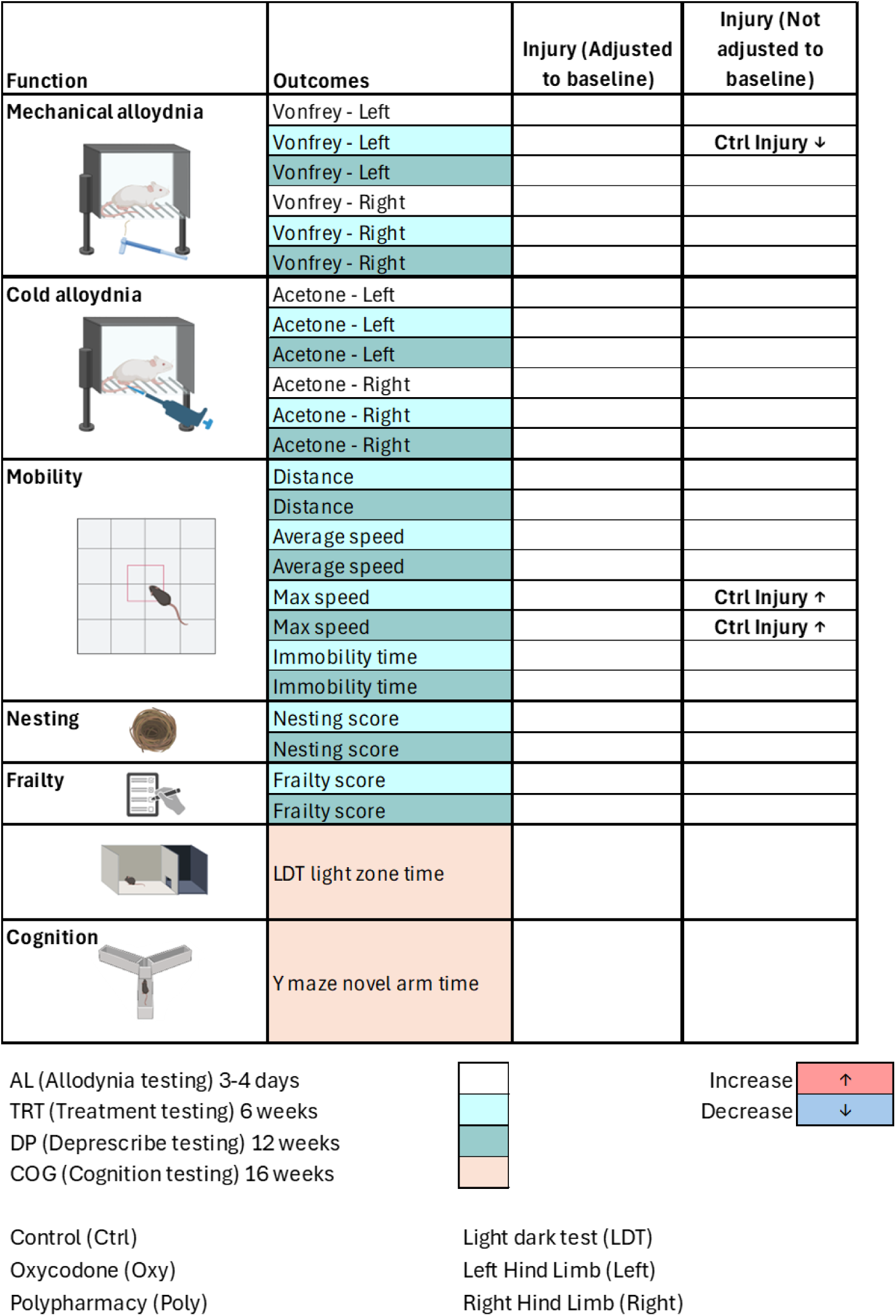
After adjusting for baseline, no injury effect was observed comparing control no injury (Sham) with control OA injured male and female mice. Summary table of outcomes shows adjusted and non-adjusted results. The following colours represent change in Injury effect over the durations (post – pre); Allodynia testing (White); 3 or 4 days of treatment and sham/injury, Treatment testing (Light blue); 6 weeks, Deprescribing testing (turquoise): 12 weeks, Last testing of study(beige): raw data at 16 weeks. GLM significant difference as indicated by p<0.05, I: Injury main effect. No Injury*treatment or Injury*sex interaction was observed. Multiple comparisons were adjusted for with Benjamini Hochberg with FDR 0.05. Comparison of groups at baseline are shown in Supplementary Table 4.

**Supplementary figure 6.**
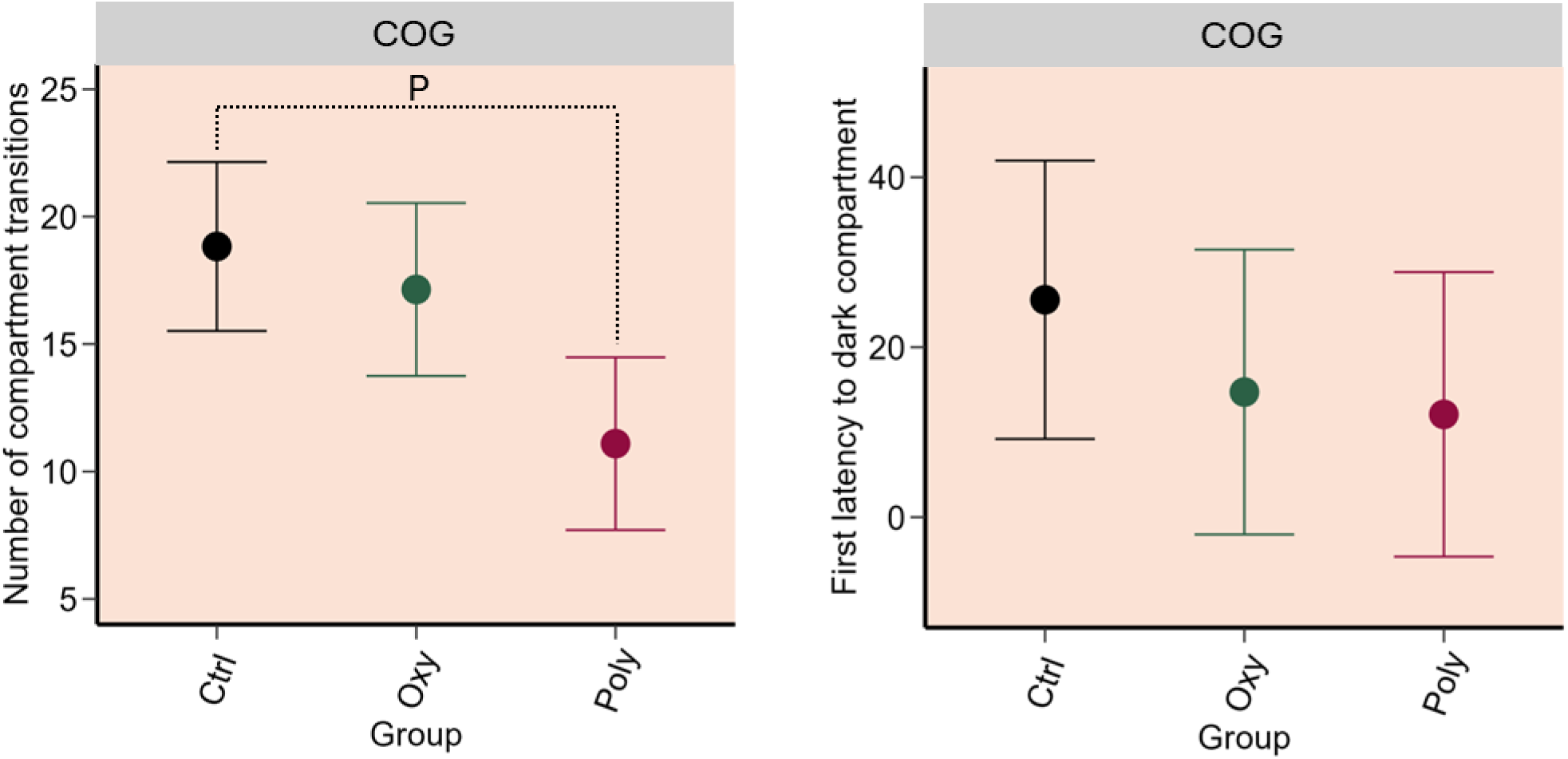
Light Dark Test: Treatment effect of control (Ctrl; black), oxycodone (oxy; green) and polypharmacy (Poly; Burgundy) on OA injured male and female mice at Cognition testing point (COG; beige, raw data at 16 weeks from OA). (a) Number of compartment transitions, (b) first latency to dark compartment. Data displayed as mean ± 95% Confidence intervals. Significant difference as indicated by p<0.05 via GLM, For all durations; O: Ctrl vs Oxy, P: Ctrl vs Poly, γ: Oxy vs Poly. No sex*treatment interaction was observed. Multiple comparisons were adjusted for with Benjamini Hochberg with FDR 0.05.

**Supplementary figure 7.**
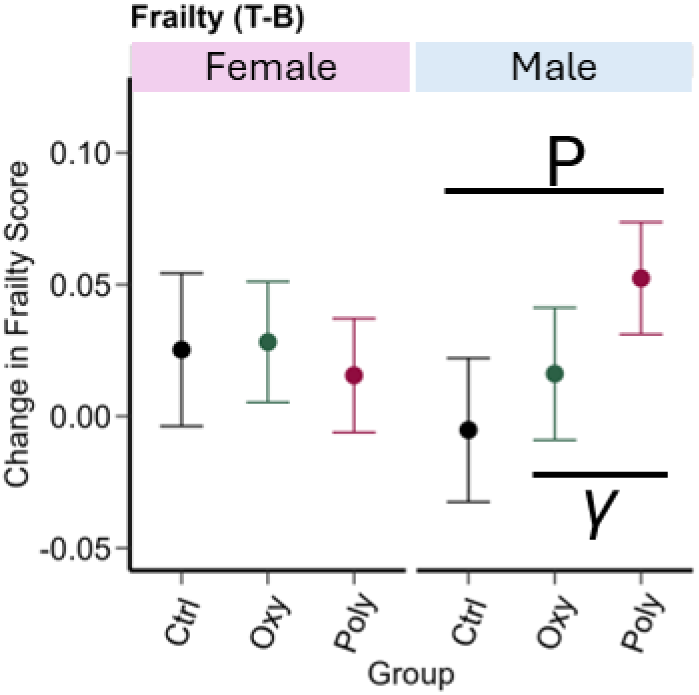
Sex differences with frailty following chronic treatment effect of control (Ctrl; black), oxycodone (oxy; green) and polypharmacy (Poly; Burgundy) on OA injured male and female mice. The graphs represent change in treatment effect over the durations of treatment (Treatment (T) – Baseline (B) time point) for Frailty score. Data displayed as mean ± 95% Confidence intervals. Significant difference as indicated by p<0.05 via mixed model with repeated measures applying a first-order heterogenous structure, LSM were generated to compare treatment groups; O: Ctrl vs Oxy, P: Ctrl vs Poly, γ: Oxy vs Poly. Multiple comparisons were adjusted with Benjamini Hochberg with FDR 0.05.

**Supplementary figure 8.**
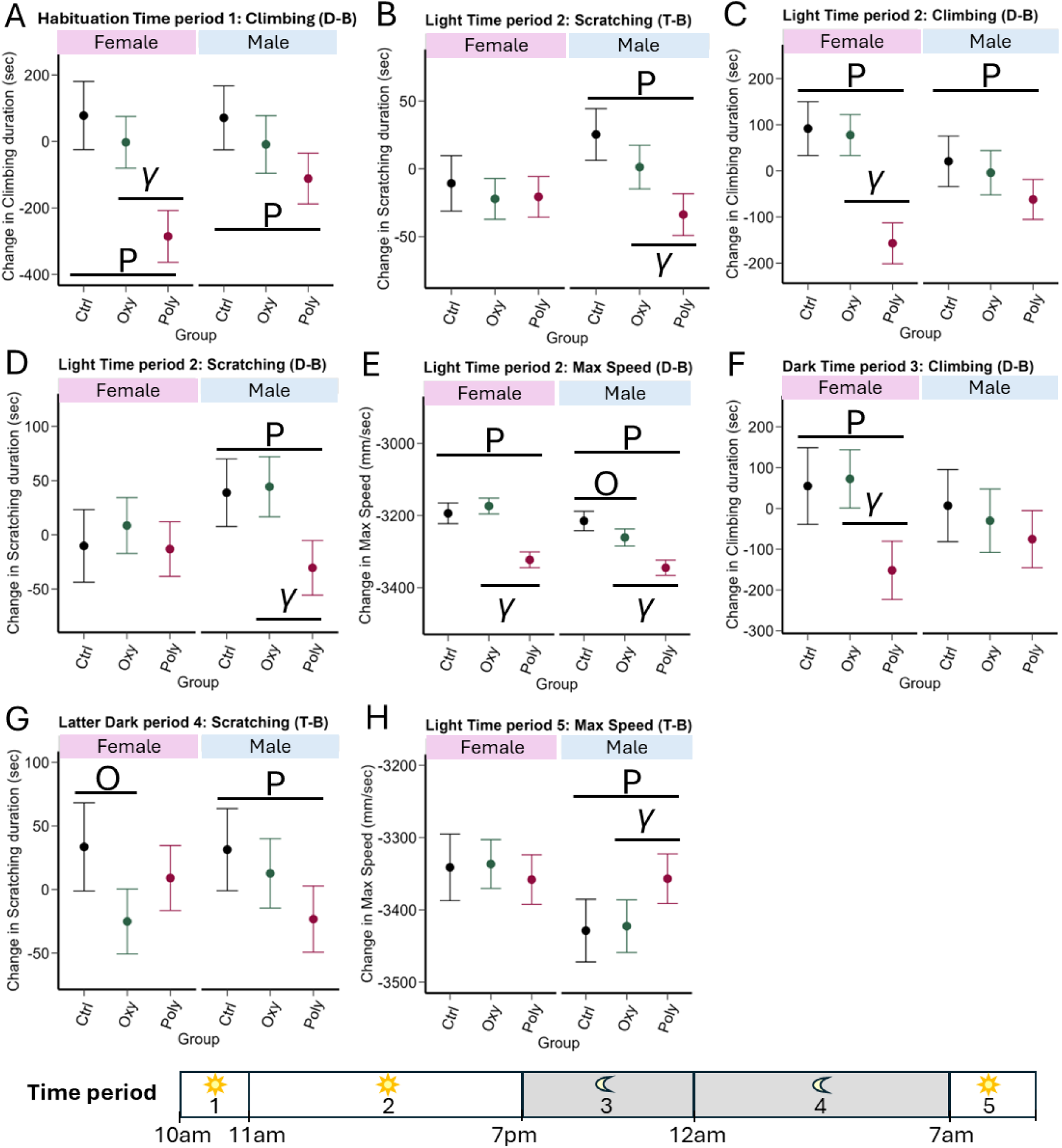
Sex differences with chronic treatment effect of control (Ctrl; black), oxycodone (oxy; green) and polypharmacy (Poly; Burgundy) on OA injured male and female mice detected using the LABORAS. The graphs represent change in intervention effect over the durations (post – pre; Time points; B: Baseline, T: Treatment and D: Deprescribe). At Habituation period 1: 10am-11am, for durations in (A) Climbing (D-B). At light period 2: 11am-7pm, for durations in (B) scratching (T-B), (C) Climbing (D-B), (D) Scratching (D-B) and (E) maximum speed (D-B). At first dark period 3: 7pm-12am, for durations in (F) Climbing (D-B). At Latter dark period 4:12am-7am, for durations in (G) scratching (T-B). At last light period 4:7am-9am, for (H) maximum speed (T-B). Data displayed as mean ± 95% Confidence intervals. Significant difference as indicated by p<0.05 via mixed model with repeated measures applying a first-order heterogenous structure, LSM were generated to compare treatment groups; O: Ctrl vs Oxy, P: Ctrl vs Poly, γ: Oxy vs Poly. Multiple comparisons were adjusted with Benjamini Hochberg with FDR 0.05.

**Supplementary figure 9.**
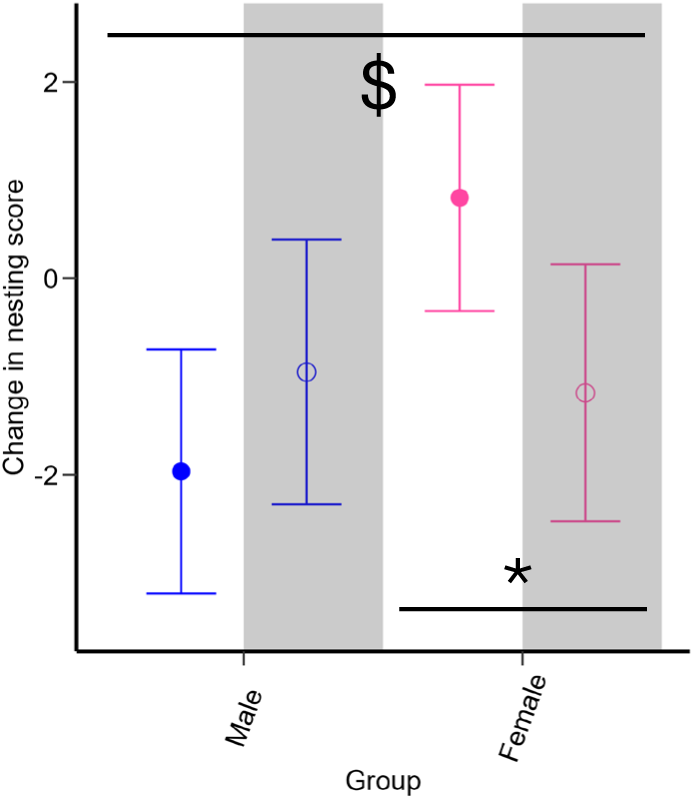
Sex impact on deprescribing oxycodone on OA injured male and female mice treated with oxycodone or polypharmacy. Graph of outcome; nesting score. Deprescribing testing (white) compares post (12 weeks) with pre (6 weeks). Males are shown in blue lines and Females in pink lines. Deprescribing group is denoted by grey shade and open circle bars . Data displayed as raw mean ± 95% Confidence intervals. Significant difference as indicated by p<0.05, δ: deprescribing main effect, # treatment*deprescribing interaction, $ sex*deprescribing interaction, *: p<0.05 as comparing corresponding treatment groups shown by the bar. LS means was conducted to compare treatment groups. Multiple comparisons were adjusted for with Benjamini Hochberg with FDR 0.05.

